# Pathway Polygenic Risk Scores (pPRS) for the Analysis of Gene-environment Interaction

**DOI:** 10.1101/2024.12.16.628610

**Authors:** W. James Gauderman, Yubo Fu, Bryan Queme, Eric Kawaguchi, Yinqiao Wang, John Morrison, Hermann Brenner, Andrew Chan, Stephen B. Gruber, Temitope Keku, Li Li, Victor Moreno, Andrew J Pellatt, Ulrike Peters, N. Jewel Samadder, Stephanie L. Schmit, Cornelia M. Ulrich, Caroline Um, Anna Wu, Juan Pablo Lewinger, David A. Drew, Huaiyu Mi

## Abstract

A polygenic risk score (PRS) is used to quantify the combined disease risk of many genetic variants. For complex human traits there is interest in determining whether the PRS modifies, i.e. interacts with, important environmental (E) risk factors. Detection of a PRS by environment (PRS x E) interaction may provide clues to underlying biology and can be useful in developing targeted prevention strategies for modifiable risk factors. The standard PRS may include a subset of variants that interact with E but a much larger subset of variants that affect disease without regard to E. This latter subset will ‘water down’ the underlying signal in former subset, leading to reduced power to detect PRS x E interaction. We explore the use of pathway-defined PRS (pPRS) scores, using state of the art tools to annotate subsets of variants to genomic pathways. We demonstrate via simulation that testing targeted pPRS x E interaction can yield substantially greater power than testing overall PRS x E interaction. We also analyze a large study (N=78,253) of colorectal cancer (CRC) where E = non-steroidal anti-inflammatory drugs (NSAIDs), a well-established protective exposure. While no evidence of overall PRS x NSAIDs interaction (p=0.41) is observed, a significant pPRS x NSAIDs interaction (p=0.0003) is identified based on SNPs within the TGF-β / gonadotropin releasing hormone receptor (GRHR) pathway. NSAIDS is protective (OR=0.84) for those at the 5^th^ percentile of the TGF-β/GRHR pPRS (low genetic risk, OR), but significantly more protective (OR=0.70) for those at the 95^th^ percentile (high genetic risk). From a biological perspective, this suggests that NSAIDs may act to reduce CRC risk specifically through genes in these pathways. From a population health perspective, our result suggests that focusing on genes within these pathways may be effective at identifying those for whom NSAIDs-based CRC-prevention efforts may be most effective.

**Author Summary:** The identification of polygenic risk score (PRS) by environment (PRSxE) interactions may provide clues to underlying biology and facilitate targeted disease prevention strategies. The standard approach to computing a PRS likely includes many variants that affect disease without regard to E, reducing power to detect PRS x E interactions. We utilize gene annotation tools to develop pathway-based PRS (pPRS) scores and show by simulation studies that testing pPRS x E interaction can yield substantially greater power than testing PRS x E, while also integrating biological knowledge into the analysis. We apply our method to a large study of colorectal cancer to identify a significant pPRS x NSAIDs interaction (p=0.0003) based on SNPs within the TGF-β / gonadotropin releasing hormone receptor (GRHR) pathway. Our findings suggest that focusing on genetic susceptibility within biologically informed pathways may be more sensitive for identifying exposures that can be considered as part of a precision prevention approach.

## Introduction

Gene-environment (GxE) interactions likely play an important role in the etiology of most complex human traits^1^. A GxE analysis aims to identify genetically defined subsets of the population that may be more sensitive to adverse or protective effects of an exposure on disease risk. Alternatively, one can view G x E interaction as investigating whether a particular exposure stimulates or suppresses the effect of a gene on disease risk. The power to detect GxE interactions, particularly in the context of a genomewide scan, is lower than the power to detect similarly-sized genetic or environmental main effects^2^. Identification of actionable GxE interactions is essential to precision medicine approaches that are expected to transform the future of medicine, particularly for primary prevention of diseases.

A polygenic risk score (PRS) is commonly used to summarize the overall effect of a collection of identified genetic variants on a particular trait. The variants used to construct the PRS can be focused on a relatively small set identified by a prior GWAS or a much larger set that captures genome-wide genetic variation. The PRS can be used to characterize the total trait variance attributable to discovered variants or to identify specific subsets of the population likely to be at highest risk for disease^3^. PRS can also be used in Mendelian randomization analysis when the disease risk factor of interest is itself predictable based on prior GWAS-discovered variants^4^.

Recently, many investigators have utilized PRS x E analysis to study gene-environment interactions for a wide range of traits, including lung cancer^5^, diabetes^6^, ADHD^7^, and cardiovascular disease^8^. Compared to single-variant GxE analysis, PRS x E analysis may provide increased power because it focuses on known disease-related variants and it integrates the signals across those variants into a potentially more informative single measure of genetic susceptibility^9^. Detecting a PRS x E interaction will allow us to answer questions such as: Does the effect of a particular exposure on disease risk vary depending on overall genetic susceptibility? Do we need to consider specific exposures when making PRS-based risk predictions? Is there a particularly high-risk subgroup, defined by both genetic susceptibility and exposure, for whom targeted prevention (e.g. early screening) may be indicated?

Despite these advantages, a potential difficulty in identifying PRS x E is that standard construction of the PRS includes all GWAS-significant variants or a very large set of genomewide variants. Environmental factors likely work to affect disease risk by altering the functioning or expression of genes within specific pathways. Examples include smoking affecting DNA repair pathways to alter lung cancer risk^10^ and red meat affecting inflammatory response pathways to affect colorectal cancer risk^11^. While a standard PRS may include several variants within an exposure-relevant pathway, its standard construction will tend to ‘water down’ the specific signals most important for identifying the interaction(s).

To overcome this challenge, we propose the use of pathway polygenic risk scores (pPRS) in gene-environment interaction analyses. Relative to a PRS, a pPRS may include a greater proportion of disease-related SNPs that individually or in combination interact with a particular exposure, and which in turn should provide greater power for detecting pPRS x E compared to PRS x E. We will describe the use of available functional annotation databases to define subsets of PRS SNPs according to their known pathway affiliation. Multiple pPRS can be constructed, each corresponding to a particular pathway and utilizing a subset of the overall collection of PRS SNPs. The use of pathway-specific PRS has been described for classifying disease subtypes^12–14^ and enhancing drug target discovery^15^, but to our knowledge not for identifying pPRS x E interactions. To illustrate our approach, we analyze PRS x E and pPRS x E interactions in a large study of colorectal cancer, focusing on over 200 GWAS-identified SNPs and a well-established protective exposure, non-steroidal anti-inflammatory drug (NSAID) use.

## Results

### Simulations

We designed a simulation study to determine whether power to detect pPRS x E interaction may be higher than for PRS x E interaction, and if so, under what conditions one may expect greater power. Briefly, we simulated 1,000 SNPs, of which 20 were assumed to affect disease (D) risk and 980 to have no effect on D. We also simulated a binary exposure (E) and generated 5 of the 20 SNPs to also have a GxE effect on D. We assumed 5 of the 1,000 SNPs fell within a pathway and varied how many of those 5 pathway SNPs overlapped with the 5 GxE SNPs, the 15 other disease-causing SNPs, and the remaining 980 null SNPs. We replicated the simulation 1,000 times and estimated power based on the proportion of replicates in which we detected interaction based on analysis of PRS x E vs. pPRS x E. Additional details of the simulation design, as well as demonstration that Type I error is preserved, are provided in Materials and Methods.

Across a wide range of simulated scenarios, power to detect interaction is greater for pPRSxE than for PRSxE (Table 1). With 20 simulated disease-causing SNPs, there was a cross-replicate average of 18.2 SNPs identified by GWAS and used for constructing the overall PRS, including an average of 4.7 of those 5 SNPs simulated to have a GxE interaction. Power to detect PRSxE interaction using the overall PRS ranged between 41% and 45% across multiple scenarios. When the 5 SNPs simulated to have a GxE effect were synonymous with the 5 SNPs in the pathway, power of the pPRSxE test was substantially higher (90%, scenario 1). This demonstrates the increased efficiency in focusing on a well-chosen subset of SNPs and corresponding pPRSxE test rather than attenuating the interaction signal in an overall PRSxE test.

**Table 1:** Power to detect polygenic risk score by E interactions.

| Sim | # Pathway SNPs | Pathway-SNP Effects on D |  |  | Power |  |  |
| --- | --- | --- | --- | --- | --- | --- | --- |
|  |  | GxE - D | G - D only | No effect | PRS x E | pPRS x E | npPRS x E |
| 1 | 5 | 5 | 0 | 0 | 44% | 90% | 2% |
| 2 | 5 | 4 | 1 | 0 | 41% | 74% | 4% |
| 3 | 5 | 3 | 2 | 0 | 41% | 47% | 7% |
| 4 | 5 | 2 | 3 | 0 | 45% | 23% | 17% |
| 5 | 5 | 1 | 4 | 0 | 45% | 7% | 28% |
| 6 | 5 | 4 | 0 | 1 | 41% | 84% | 4% |
| 7 | 5 | 3 | 0 | 2 | 41% | 69% | 7% |
| 8 | 5 | 2 | 0 | 3 | 45% | 52% | 14% |
| 9 | 5 | 1 | 0 | 4 | 45% | 27% | 25% |

Simulated power based on 1,000 replicates. Each replicate includes 15 SNPs with a G-only effect on D and 5 SNPs with a GxE effect on D. There are 5 SNPs in the pathway. Each simulation scenario varies the number of pathway SNPs that overlap with the GxE SNPs (GxE-D), G-only SNPs (G-D), and no-effect SNPs. Power is the proportion of replicates in which the null hypothesis of no interaction is rejected when the polygenic score is based on all GWAS significant SNPs (PRS x E), GWAS SNPs in the pathway (pPRS x E) or GWAS SNPs not in the pathway (npPRS x E).

We also considered simulation scenarios in which only a subset of the 5 pathway SNPs overlapped with the 5 GxE SNPs. These included scenarios in which the pathway SNPs without a GxE effect either did (Table 1, Scenarios 2-5) or did not (Scenarios 6-9) have a main (G only) effect on the trait. When the 5 pathway SNPs include 4 with true GxE and 1 G-only (scenario 2) or 3 GxE and 2 G-only (scenario 3), power of the pPRSxE test was still greater (74%, 47%, respectively) than the PRSxE test. However, with 2 GxE and 3 G-only (scenario 4) or 1 GxE and 4 G-only (scenario 5), power of the pPRSxE was lower (23%, 7%, respectively). By comparison, when the 5 pathway SNPs included 4 with true GxE and 1 with no effect on the trait (scenario 6), power was 84%, larger than the 74% when the non-GxE SNP had a G-only effect (scenario 2). This is because in scenario 6 the non-GxE SNP likely is not discovered in the initial GWAS and thus is not used in forming the pPRS (or PRS) score, and therefore is not attenuating the signal in the remaining GxE SNPs. This trend is further exemplified by the corresponding higher powers in scenarios 7, 8, and 9 compared to scenarios 3, 4, and 5, respectively.

### Colorectal Cancer (CRC) Application

The most recent and largest GWAS of CRC described a total of 204 previously identified and novel autosomal SNPs that reached genome-wide significance^16^. We investigated whether PRS and pPRS formed from these SNPs interact with use of aspirin or non-steroidal anti-inflammatory drugs (NSAIDs) use, a factor well-established to reduce CRC risk^17–19^. We used data from the Functionally Informed Gene-environment Interaction (FIGI) study, a consortium of 45 studies that includes 78,253 subjects (33,937 cases, 44,316 controls) with complete data on NSAIDs, genotypes, and covariate data^19^. Adjusting for covariates, the NSAIDs main effect on CRC is OR=0.76 (95% C.I. 0.74, 0.79). Although NSAIDs is a protective factor on average, there are risks associated with regular use, such as gastrointestinal bleeding, that necessitate a precision prevention approach. This is one motivation for exploring a precision prevention approach for NSAIDs based on possible modification by genetic susceptibility.

We constructed an overall PRS by first applying logistic regression within the FIGI sample to model CRC as a function of the 204 GWAS SNPs, with adjustment for study, sex, age, and three global ancestry PCs (see Materials and Methods). The SNP-specific log-odds ratios estimated from this model were used as the weights *[w]* to construct a PRS_i_, i=1, …, N for each study subject. To construct pPRS, we first used ANNOQ^20^ which successfully annotated 189 of the 204 SNPs to 265 protein-coding genes. The remaining 15 SNPs were mapped to non-coding genes and are ignored in this analysis. Application of PANTHER^21^ annotated 66 of the 265 genes to a total of 50 pathways (Figure 1), with pathways for the remaining 199 genes not identified. Among the 50 pathways, four of them included more genes than expected by chance alone at a false discovery rate (FDR) of 5%, identified by a Fisher’s Exact test in PANTHER (Table 2). These included the TGF-β signaling pathway (raw p=5.8×10^−6^), Alzheimer disease presenilin pathway (p=5.8×10^−5^), Gonadotropin-releasing hormone receptor pathway (p=4.8×10^−5^), and the Cadherin signaling pathway (p=1.38×10^−3^). A total of 30 of the 204 SNPs were annotated to genes in these pathways. Subsets of the above PRS weights were utilized to construct the corresponding four pPRS scores. Annotated genes in the TGF-β signaling (TGF-β) pathway and Gonadotropin-releasing hormone receptor (GRHR) pathways are highly overlapped (Figure 2A), as are genes in the Cadherin signaling (CADH) and Alzheimer’s disease presenilin(ALZ) pathways (Figure 2B). These overlaps lead to significant correlations between the computed pPRS scores for TGF-β and GRHR (R^2^=0.58) and for CADH and ALZ (R^2^=0.71). Given this, we also constructed two additional pPRS scores based on SNPs within the combined subsets of TGF-β/GRHR genes and CADH/ALZ genes, respectively.

**Figure 1:**
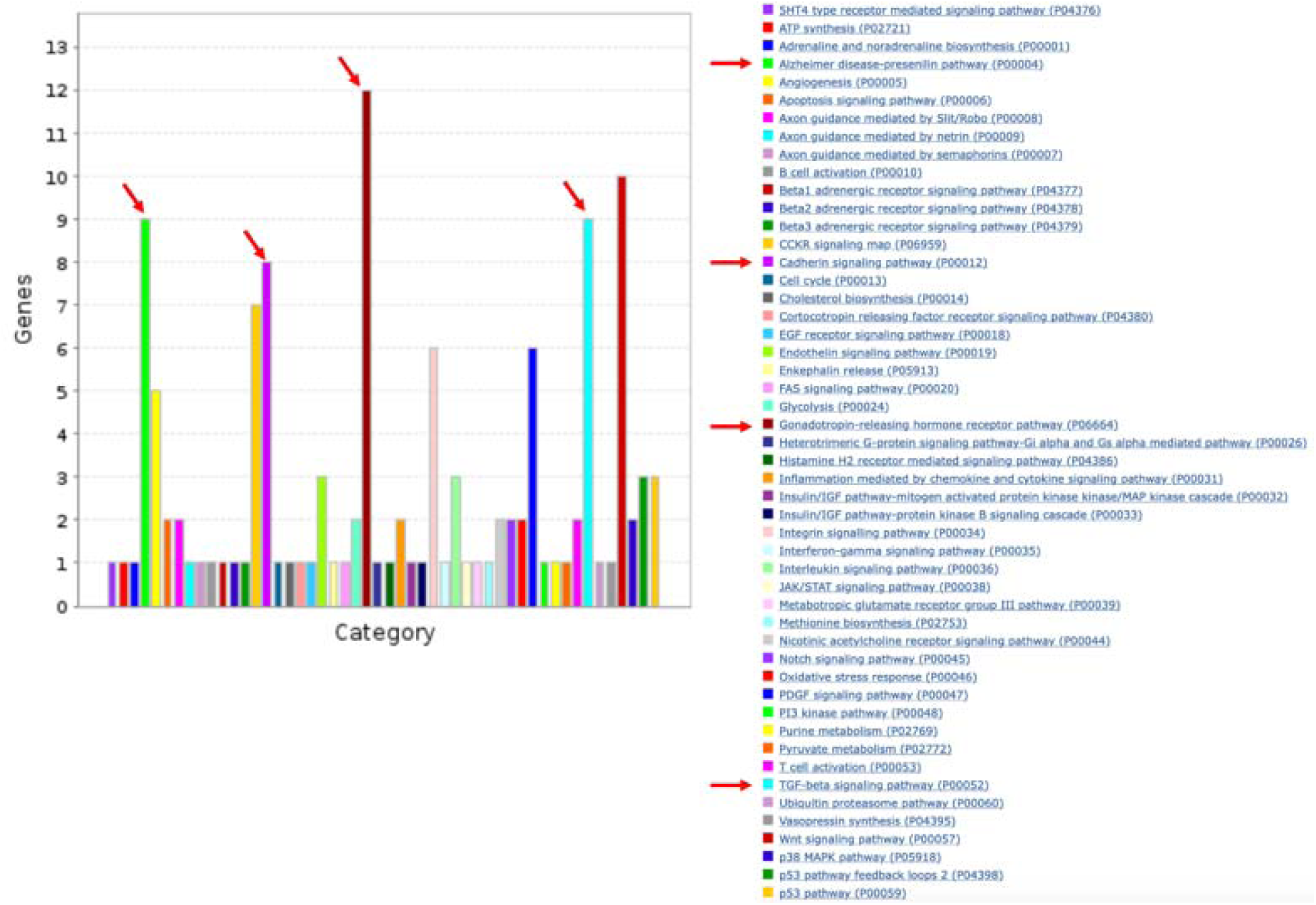
Number of genes and pathways represented among the 204 CRC-associated SNPs. Arrows indicate the four pathways with significant over-representation compared to expected (Table 2).

**Table 2:** Four pathways significantly over-represented among the 204 CRC-relate SNPs.

|  | Homo sapiens (REF) | AnnoQ_uniq_genes |  |  |  |  |  |
| --- | --- | --- | --- | --- | --- | --- | --- |
| PANTHER Pathways | # | # | expected | Fold Enrichment | +/- | raw P value | FDR |
| TGF-beta signaling pathway | 100 | 9 | 1.29 | 6.99 | + | 5.82E-06 | 4.66E-04 |
| Alzheimer disease-presenilin pathway | 127 | 9 | 1.64 | 5.50 | + | 4.01E-05 | 2.14E-03 |
| Gonadotropin-releasing hormone receptor pathway | 231 | 12 | 2.97 | 4.03 | + | 4.83E-05 | 1.93E-03 |
| Cadherin signaling pathway | 163 | 8 | 2.10 | 3.81 | + | 1.27E-03 | 4.06E-02 |

Column 2 shows the total number of genes in Homo sapiens annotated to the indicated pathway.
Column 3 shows the number of genes from the pathway that are among the genes linked by application of AnnoQ to the 204-CRC associated SNPs

**Figure 2:**
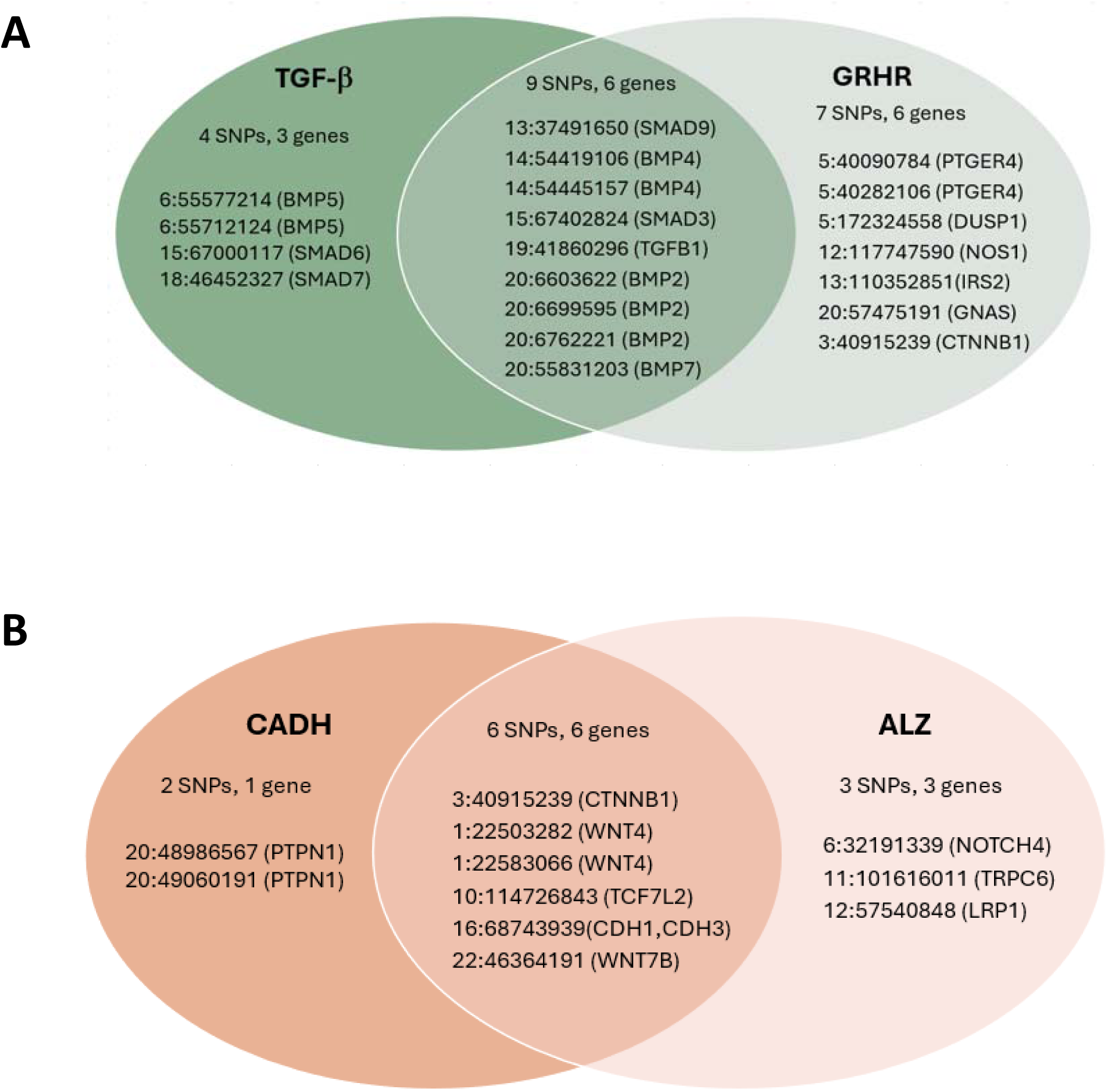
A. Subset of 204 CRC-associated SNPs annotated to genes within the TGF-β and/or the Gonadotropin releasing hormone receptor (GRHR) pathways. **B.** Subset of 204 CRC-associated SNPs annotated to genes within the Cadherin signaling (CADH) and/or the Alzheimer’s disease-presenilin (ALZ) pathways.

The estimated GxE odds ratio (OR_GxE_) for the overall PRS x NSAIDs interaction is 0.99 and is not statistically significant (p=0.41, Table 3). We also did not observe significant pPRS x E interactions for the CADH and ALZ pathways. However, the pPRS x NSAIDs interaction was significant for both the TGF-β (OR_GxE_=0.96, p=0.0069) and GRHR (OR_GxE_=0.96, p=0.016) pathways. The TGF-β and GRHR pathways combined include 20 of the 204 SNPs (Figure 2A). The pPRS x NSAIDs interaction is more pronounced (OR_GxE_=0.94, p=0.0003) based on the pPRS formed from this joint set of TGF-β and GRHR SNPs (Table 3). This estimate can be interpreted as an additional 0.94 protective effect of NSAIDs on CRC risk per increase of 1 standard deviation in the combined TGF-β/GRHR pPRS. To further explore this interaction, we used the model to predict the NSAIDs effect on CRC at various percentiles of the TGF-β/GRHR pPRS (Figure 3). For those at the 5^th^ percentile of the pPRS (low risk), the estimated NSAIDs OR is 0.84 (0.79, 0.89) while at the 95^th^ percentile (high risk), it is 0.70 (0.65, 0.74). Put another way, regular NSAIDs use is predicted to reduce CRC risk by 16% for those at low risk based on the TGF-β/GRHR pPRS and by 30% for those at high TGF-β/GRHR pPRS risk.

**Table 3:** Analysis of polygenic risk score x NSAIDs interactions for Colorectal Cancer.

| PRS Type | PRS |  | E (NSAIDs use) |  | PRS x E |  | p-value<sup>b</sup> |
| --- | --- | --- | --- | --- | --- | --- | --- |
|  | OR<sup>a</sup> | (95% CI) | OR | (95% CI) | OR | (95% CI) |  |
| PRS: All SNPs<sup>*</sup> | 1.63 | (1.61, 1.66) | 0.76 | (0.74, 0.79) | 0.99 | (0.95, 1.02) | 0.41 |
| <u>4 Pathways<sup>&</sup></u> |  |  |  |  |  |  |  |
| pPRS: TGF-β | 1.18 | (1.16, 1.20) | 0.76 | (0.74, 0.79) | **0.96** | **(0.93, 0.99)** | **0.0069** |
| pPRS: Gonadotropin receptor | 1.17 | (1.15, 1.19) | 0.76 | (0.74, 0.79) | **0.96** | **(0.93, 0.99)** | **0.016** |
| pPRS: Cadherin signaling | 1.10 | (1.09, 1.12) | 0.76 | (0.74, 0.79) | 1.00 | (0.97, 1.04) | 0.82 |
| pPRS: Alzheimer's presenillin | 1.09 | (1.08, 1.11) | 0.76 | (0.74, 0.79) | 0.99 | (0.96, 1.02) | 0.46 |
| <u>2 combined Pathways<sup>&</sup></u> |  |  |  |  |  |  |  |
| pPRS: TGF-β and/or Gonadotropin receptor | 1.21 | (1.19, 1.23) | 0.76 | (0.74, 0.79) | **0.94** | **(0.92, 0.97)** | **0.0003** |
| pPRS: Cadherin and/or Alzheimer's presenillin | 1.11 | (1.10, 1.13) | 0.76 | (0.74, 0.79) | 1.00 | (0.97, 1.03) | 0.86 |
| PRS Other<sup>#</sup> | 1.55 | (1.53, 1.58) | 0.76 | (0.74, 0.79) | 1.01 | (0.98, 1.04) | 0.63 |

<sup>\*</sup> PRS formed based on 204 GWAS significant SNPs as reported in Fernandez-Rozadilla et al. (2022)
<sup>&</sup> pPRS based on subsets of the 204 SNPs within the indicated pathway
<sup>#</sup> PRS based on the subset of 174 of the 204 SNPs that are not within any of the indicated pathways
<sup>a</sup> Odds ratios (OR) are scaled to a 1 s.d. increase for the indicated PRS and compare users to non-users for NSAIDs. All $p < 10^{-10}$ .
<sup>b</sup> p-value for the test of the null hypothesis of no PRS x E interaction.

**Figure 3:**
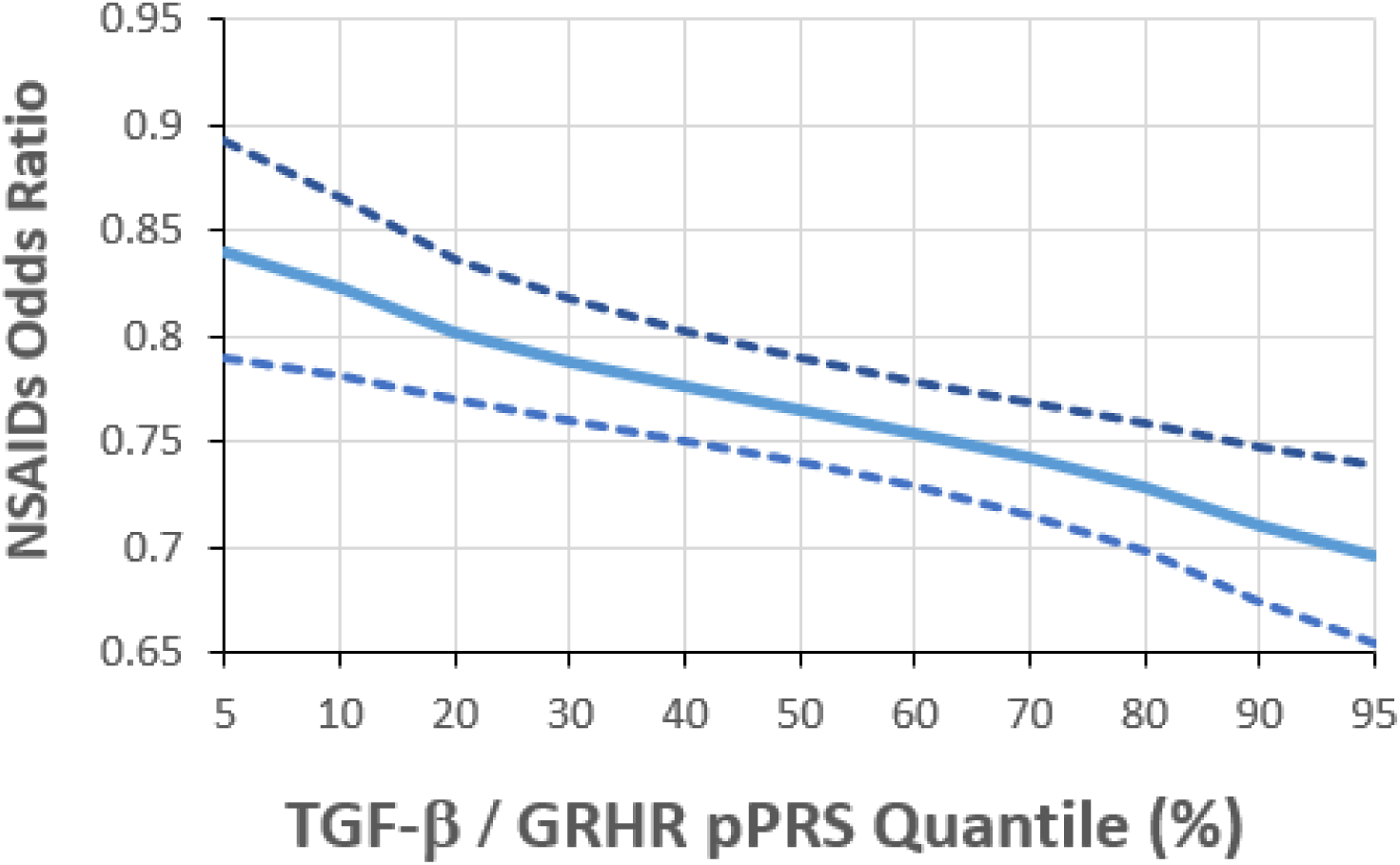
NSAIDs odds ratio for CRC by quantile of the joint TGF-β / GRHR pPRS quantile.

We repeated these analyses utilizing PRS weights obtained from the PGS catalog for the same set of SNPs (PGS-ID 003850). This was done to further evaluate how use of our own data to estimate PRS weights (as above) compared to the more standard approach of using catalog-derived, published weights. Applying the two sets of weights to our analysis sample yielded PRS scores that were very highly correlated for the overall PRS (R^2^=0.9) as well as for the TGF-β (0.98), GRHR (0.97), CADH (0.97), and ALZ (0.89) pPRS. Not surprisingly, then, results based on PGS catalog weights (Table 4) were very similar to those reported above (Table 3), with similar interaction estimates and levels of significance for TGF-β, GRHR and the joint TGF-β/GRHR pPRS x NSAIDs effects, and non-significant results for the other pathway and overall PRS x NSAIDs tests.

**Table 4:**
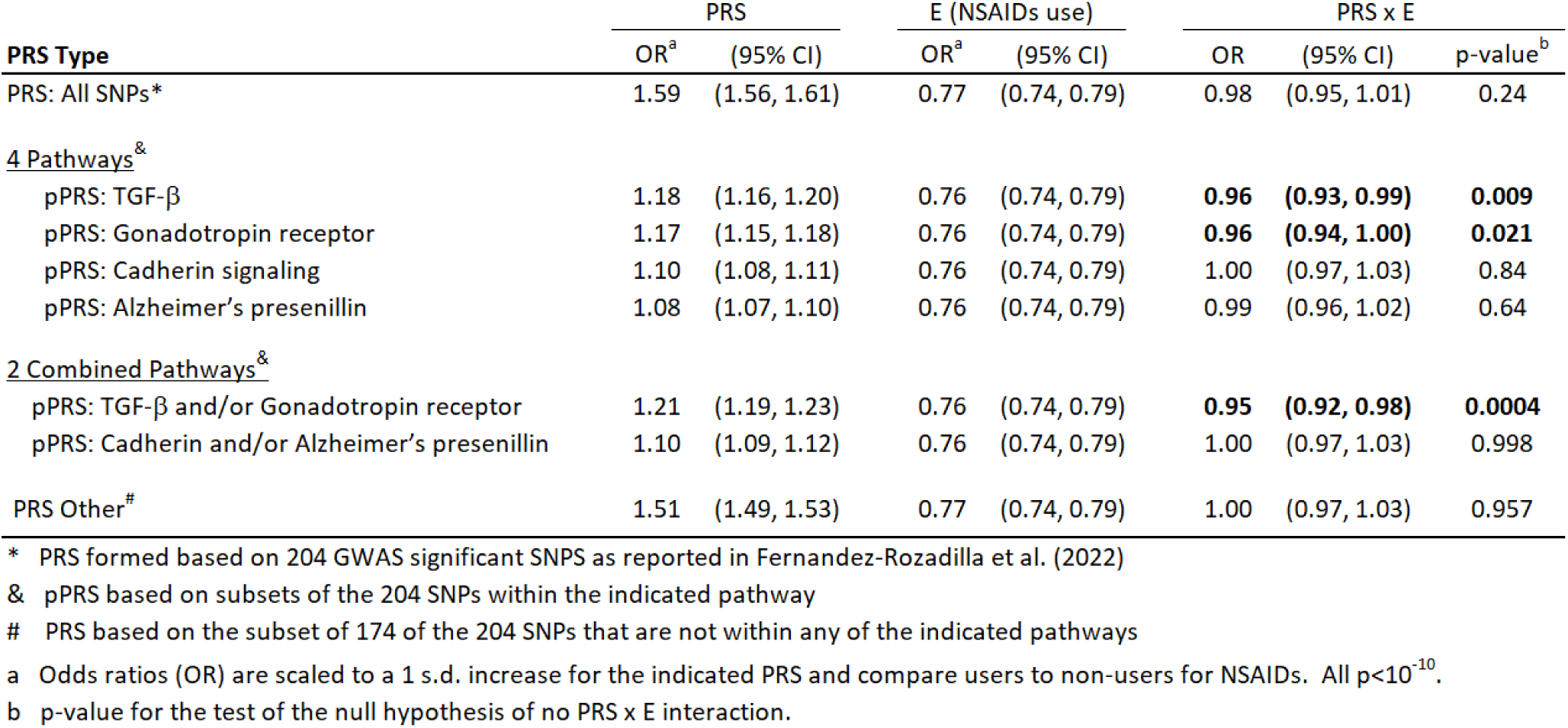
Analysis of PGS Catalog derived polygenic risk score x NSAIDs interactions for Colorectal Cancer.

## Discussion

We have demonstrated by simulation and application to data that forming a PRS based only on a subset of GWAS significant SNPs, specifically a subset defined a priori based on pathway information, has the potential to better identify novel PRS x E interactions. We also demonstrate that power may be reduced using the standard practice of testing PRS x E interaction based only on an overall PRS. This power reduction is likely due to ‘watering down’ the interaction signal with the inclusion of most of the SNPs in the PRS construction that do not have any role in modifying the effect of E on disease. By contrast, the use of external pathway information to form a pPRS has the potential to improve power by focusing on genetic variation within a particular pathway that modifies the E effect. Examination of E effects across quantiles of the pPRS can identify those genetically-defined subsets that are most affected, or protected, by exposure. For example, our analysis of CRC suggests that although NSAIDs use is generally beneficial for all, those with the highest TGF-β/GRHR pathway PRS experience a significantly greater reduction in CRC relative risk with regular NSAIDs use. This result both adds to the overall preventive evidence for NSAIDs on CRC risk and suggests possible biological pathways that are involved in this action.

The use of pPRS in interaction testing relies on external information to identify the pathways corresponding to a particular set of SNPs. In this paper, we focused on the set of GWAS significant SNPs, but one could relax the criteria to include SNPs that have GWAS p-value that achieves a lesser threshold (e.g. 5×10^−5^) or to a wider collection without regard to GWAS significance. As has been previously shown, a GxE interaction typically induces a direct disease-gene (DG) association^22–25^, and so requiring some level of DG association to be included in PRSxE or pPRSxE analysis is reasonable. In our application to CRC, we created a workflow that utilized AnnoQ to annotate SNPs to genes and PANTHER to annotate genes to pathways. We recognize that there are alternative tools/databases that could be employed, for example Reactome (reactome.org) or Gene Ontology (geneontology.org), and that different workflows would likely result in pathway assignments that do not fully overlap. A particular application of pPRS x E analysis could consider the use of multiple workflows, each using different tools/databases, to evaluate the sensitivity of findings to specific pathway definitions and corresponding SNP/gene assignments.

An ancillary finding in this paper is the demonstration that one can construct a PRS or pPRS in three different ways if the ultimate focus is a valid test of interaction. Approach #1 (Methods), i.e. to obtain existing PRS weights from the PGS catalog, is the one most often used. This has the advantages that the weights are typically estimated using a large and independent dataset, and that one can apply the weights to their data to estimate both PRS main and interactive effects. A potential disadvantage, however, is that the data used to generate the PGS weights may come from a population(s) that do not represent the sample used for PRS x E analysis. It is well known that cross-population application of PRS for main effects can lead to poor estimation, and the same will hold for analysis of PRS x E interactions. The advantage of Approach #2 (Methods) is that it leverages the discovery of SNPs in a larger, independent population, but tailors the weights used in PRS construction to the specific population being studied for interaction. Of course, this is also not free of cross-population issues if the discovered SNPs in the independent population are not representative of the SNPs/genes affecting the trait in the study population. Approach #3, in which the study sample is used to discover SNPs and estimate weights, is perhaps the cleanest from the standpoint of population heterogeneity but may suffer from reduced power to discover SNPs relative to larger independent studies. As we demonstrated in our CRC analysis, the flexibility to use any of these approaches for valid interaction testing provides the opportunity to evaluate the robustness of PRSxE and/or pPRSxE findings to the choice of PRS SNPs and weights.

Our results highlight that pPRSxE can identify pathways with functional relevance to the exposure’s putative mechanisms of action. In this case, we provide evidence that the protective effect of NSAIDs on CRC risk is modified by variation in the TGF-β and GRHR pathways. While aspirin and other NSAIDs primary inhibitory activity on PTGS1/2 (or COX1/2) have long been hypothesized as central mechanisms of for their anticancer effects, the overall mode of action is still not yet clear. Several lines of functional evidence have supported a role for the TGF-β superfamily in mediating aspirin/NSAIDs protective effects against CRC^26^, particularly in models of mismatch repair deficient CRC^27^. Long-term follow-up of the CAPP2 randomized, placebo-controlled trial conclusively demonstrated that aspirin is protective against CRC among patients with Lynch syndrome^28^, also known as hereditary non-polyposis colon cancer resulting from pathogenic variants within DNA mismatch repair genes, suggesting that NSAID protection may also extend to those with sporadic mismatch repair deficient tumors. TGF-β has also been demonstrated to induce *HPGD*^27^, a prostaglandin-degrading enzyme with tumor suppressor activity that works as a catabolic antagonist for PTGS-2 activity^29^. Moreover, HPGD mucosal gene expression has been demonstrated to stratify individuals that may be more likely to experience a preventive benefit from aspirin use^30^. While other TGF-β superfamily members like GDF15 have been proposed as potential markers for precision prevention of CRC with NSAIDs^18^, the role for bone morphogenetic proteins (BMPs) and SMAD family proteins in NSAID chemoprotection are less well established than they are for other agents, like metformin^31^, or other physiologic processes, like osteogenic differentiation^32,33^. Similarly, functional evidence is limited for a specific role of Gonadotropin-receptor pathway overall in NSAIDs mechanisms of action. However, of those genes included in the pPRS score, prior evidence links NSAID anti-cancer activity with β-catenin (CTNNB1^34–37^), GNAS^38^, and PTGER4^19^, the extracellular receptor for PGE_2_, the major downstream prostanoid produced by PTGS-2. Combined, these results highlight that a pPRSxE approach may identify additional network nodes with potential functional relevance for future mechanistic interrogation.

We have shown that leveraging prior GWAS results combined with biological information to construct subsets of SNPs in pPRS x E tests has the potential to improve power compared to SNPxE or overall PRSxE tests. An additional advantage of the pPRS x E analysis is that it may strengthen the evidence for a potential biological mechanism, via the involved pathway, by which E affects the outcome. Although we have focused on SNP subsets based on pathway information, we recognize there are other sources of information that could be used to create subsets. For example, subsets could be formed based on information on SNP-expression in a relevant tissue or cell type, or based on SNP associations with traits related to the trait of interest. Future research is needed to examine the robustness of pPRSxE analyses to the choice of annotation workflow, to the approach to creating subsets, and to demonstrate whether pPRS can be used to successfully identify novel gene-environment interactions for other complex traits.

## Materials and Methods

### Notation and Standard GxE Analysis

Let *D_i_* denote a disease indicator for subject *i*, *i*=1, …, N, *E_i_* an exposure of interest, and ***Z_i_*** a vector of adjustment covariates (e.g. age, sex, ancestry principal components). Assume one or more GWAS has been conducted, yielding a set ***G***=[*G_1_, G_2_, …, G_M_*] of trait associated SNPs, for example those with p<5×10^−8^ for the test of SNP vs. D association. Assume further that a case-control sample has been obtained, with complete data for *D, E, **Z***, and ***G*** n each subject. For analysis of *GxE* interaction with a single SNP, we assume logistic regression model of the form:

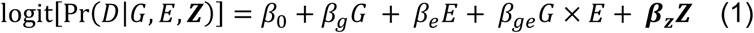

Here *β_g_* denotes the genetic ‘main’ effect quantifying the association between *G* and *D* when *E* =0, *β_e_* is the corresponding environmental main effect, and *β_ge_* parameterizes the *GxE* interaction effect of primary interest. *G* is typically coded as the number of minor alleles, 0, 1, or 2 if it is measured or the corresponding expected number if imputed. In practice, we often center both *G* and *E* on their respective sample means yielding

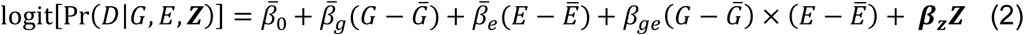

Here *β̅_g_* parameterizes the *G* to *D* association at the mean of *E* and similarly for *β̅_e_*. An advantage of this centering is that *β̅_g_* and *β̅_e_* approximate the ‘marginal’ effects of *G* and *E*, for example the direct effect of *G* on *D* (*γ_g_*) that is obtained in a GWAS using the model:

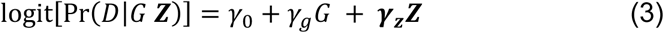

The test of *GxE* interaction evaluates the null hypothesis H_0_: *β_ge_*= 0 and can be based on a Wald, Score, or likelihood-ratio test from either model 1 or model 2, with proper adjustment to the significance level to achieve the desired family-wise error rate. If each of S SNP is tested, the significance level for each SNP is α/S, i.e. based on a Bonferroni correction for S tests subject to overall significance level α.

### Formation of the PRS

For a collection of M SNPs, e.g. those previously identified as GWAS significant, the following logistic model is used to estimate the SNP effects in the context of a single joint model:

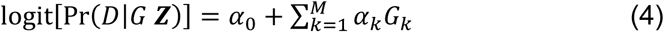

We define weights [*w_k_*] to be the estimates [*α̂*] from Model 4. The equation for generating a PRS for the i^th^ individual is

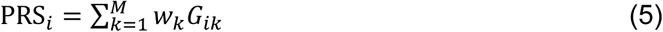

Replacing *G* in Equation 1 by the PRS yields the following model which can be used to estimate and test for PRS x E interaction:

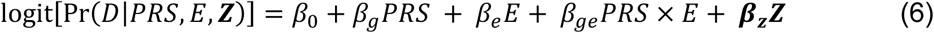

### PRS Weights

The PRS weights are typically derived from a separate resource. For example, the PGS catalog^39^ provides weights for over 650 traits, including multiple sets of weights for many of the traits. It is important that the weights come from independent data resources if the PRS will be used to examine direct risk effects on the disease of interest in the N subjects under study. In other words, if the weights are generated based on the N subjects under study, applying the resulting PRS to the same subjects will result in biased inference of the PRS effect on disease risk. However, we will demonstrate that the same dataset can be used to generate the PRS weights if the focus is on PRS x E interaction. The ability to ‘double use’ the same data to generate and apply the weights relies on the independence between the marginal genetic effects (estimated via Model 3) and the interaction effects (estimated via Model 2). This independence has been shown for tests of single SNPs^40^ and is the basis for several 2-step genomewide GxE scan methods that screen on marginal G effects in Step 1 and use the information to prioritize SNPs for GxE testing in Step 2^22,24,25,41^. We provide simulations in this paper demonstrating that this independence holds for use of the weights [*w_k_*] derived from Eq. 4 for downstream PRS x E interaction analysis.

Given this independence, there are three Approaches one might consider for generating the [*w_k_*] and corresponding PRS:

1. <u>Obtain [*w_k_*] from prior studies based on one or more independent datasets.</u> As noted above, these could come from the PGS catalog or a specific previous GWAS of the trait of interest. This will provide weights that can be applied to the N subjects under study for use in estimating PRS main and PRS x E interaction effects on D. One must be prepared to assume, however, that the weights generated from the previous population(s) are applicable to the current study population, which may not be reasonable if there are differences in ancestry^42^.
2. <u>Obtain M SNPs from prior GWAS but estimate [*w_k_*] in the current sample that will be used for PRS x E analysis</u>. Again the list of previously identified SNPs could come from the PGS catalog or a specific prior GWAS, but rather than use existing weights, model 5 is applied to the M SNPs in the current data to generate [*w_k_*]. The corresponding PRSi, i=1, …, N, would not provide valid estimates of the PRS main effect but are valid for estimating and testing PRS x E effects. An advantage of this approach is that the weights are computed based on the demographic (e.g. sex, age, ancestry) composition of the current study. The discovery of the set of M SNPs, however, may have been based on different populations with different exposure histories and thus may not fully represent the genetic and GxE contributions in the current sample.
3. <u>Conduct a GWAS on the current sample to both identify M SNPs and compute corresponding [*w_k_*]</u>. Compared to approaches 1 and 2, this has the advantage that both the selection of M SNPs and calculation of weights reflect the population structure and exposure characteristics of the current sample. On the other hand, the current sample may be smaller than prior studies and thus have less power to identify important SNPs in the GWAS discovery step.

We will demonstrate the third approach in our simulation and the first two approaches in our application to colorectal cancer.

### Pathway PRS

Human genes and their products typically function together within biological pathways to maintain proper cellular functions. SNPs located within or near gene regions have the potential to influence the pathways in which these genes are involved. We assume that the collection of M SNPs used to form the PRS include subsets of SNPs falling within different biological pathways. To assign each SNP to a pathway, we first use the Annotation Query (AnnoQ) platform^20^ to derive annotations to Ensembl^43^ and RefSeq^44^ genes using inferences from ANNOVAR^45^, SnpEff^46^ and VEP^47^. SNPs residing in enhancer regions were linked to their target genes via PEREGRINE^48^. The resulting genes were annotated to pathways using the PANTHER^21^ Classification System (v.18.0)^49^. The set of genes falling within the same pathway were tested for overrepresentation relative to the PANTHER Pathway annotation sets^50^. Each pathway that is significantly over-represented is the focus of pPRS computation and pPRS x E interaction testing.

Assuming that K pathways are identified by the above approach, we define pPRS_1_, pPRS_2_, …, pPRS_K_ to be PRS including only those SNPs within the corresponding pathway. We also let pPRS_0_ denote the PRS that includes the subset of M SNPs not annotated to any of the K pathways. Let S_k_, k=0,…,K denote the subset of M SNPs included in the k^th^ subset. The pPRS for pathway k is then defined as:

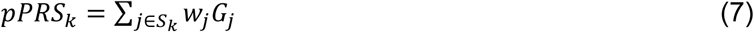

where weights are obtained by one of the three approaches described above. Note that this approach to computing pPRS implicitly assumes that the weights are generated from the full model of *D* that includes all M SNPs, which has the advantage that the weights are mutually adjusted for one another. To investigate a particular pPRS, Equation 6 can be modified to:

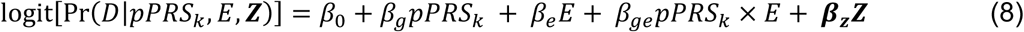

Alternatively, one can also use a model that includes all pPRS, with form:

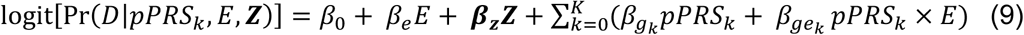

Additional interactions between pPRS and ***Z*** and/or between *E* and ***Z*** can also be included to account for potential confounding at the level of the pPRS x E effects^51^. We note that it is possible for a particular SNP to be annotated to two or more pathways. In this situation, there will be correlation between two pPRS that include the same SNP(s), which will require care in interpreting the resulting effect estimates.

### Simulation Studies

We conducted simulation studies to: 1) evaluate the claim that the same dataset can be used to estimate the PRS weights [w], construct a PRS, and obtain valid estimates and tests of PRS x E interaction, and 2) to compare the power of pPRS x E to PRS x E analysis.

We generate a dataset that includes 5,000 cases and 5,000 controls, with a binary exposure E and 1,000 randomly and independently generated SNPs per subject. We designate Q=20 of the SNPs to affect disease risk, with Q_G_ having only a main G to D effect and Q_GxE_ having both a main and GxE effect. We further assume that Q_P_ = 5 of the 1,000 SNPs fall within a particular pathway and that Q_PG_ of the pathway SNPs have only main effect and Q_PGxE_ have a GxE effect. We vary Q_PG_ and Q_PGxE_ across simulation scenarios. For each simulation scenario, we generate 1,000 replicate datasets and use these to evaluate Type I error and power We generate each G as a binary variable with 35% population prevalence and E as binary with population prevalence 50%. Conditional on simulated G and E, disease status for each subject was generated according to a random Bernoulli distribution with probability of disease (P_D_) given by:

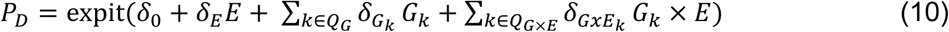

The values of 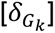 were determined using Quanto^52^ to achieve an expected power of at least 90% to detect each of the Q SNPs in a GWAS with adjustment for 1,000 tests. The 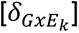 values were set to achieve approximately 10% power to detect GxE interaction for each of the Q_GxE_ SNPs, assuming 20 SNPs are evaluated for SNP x E interaction post-GWAS.

For each simulated dataset, we conducted a GWAS of the 1,000 SNPs to identify the M that were significant at the 0.05/1,000 = 5×10^−5^ level. These M SNPs were used in a model of the form in Equation 4 to generate weights [w_k_]. We computed the standard PRS based on these M weights using Equation 5, the pathway PRS (pPRS) based on Equation 7 for the subset of M within Q_P_, and the non-pathway PRS (npPRS) based on Equation 7 for the subset of M not within Q_P_. Each simulation scenario was replicated 1,000 times and we tallied the proportion of replicates in which the null hypothesis of no interaction was rejected for likelihood ratio tests of PRSxE, pPRSxE, and npPRSxE based on Equation 8. This proportion estimated Type 1 error in simulations with Q_GxE_=0 and power when Q_GxE_ > 0.

Our first set of simulations shows that use of the same data set to run a GWAS, generate PRS weights, and test PRS x E interaction (approach #3, see above) preserves the desired Type I error rate for the interaction test (Table S1). We simulate 20 disease-causing SNPs (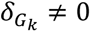 for *k* ∈ *Q_G_*) and set 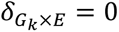, for all *k* (Eq. 11). We tested five methods to identify the SNPs to generate PRS weights: 1) Identify the M SNPs that were significant at the 0.05/1,000 = 5×10^−5^ level; 2) identify the M that were significant at the 0.05/10 = 5×10^−3^ level; 3) identify the M that were significant at the 0.05 level; 4) include the 20 disease-causing SNPs; and 5) randomly select 10 of the 20 disease-causing SNPs and 10 from the 980 null SNPs. Across all these scenarios, the estimated Type I error rate was within simulation variability of the desired 0.05 level. Since approaches #1 and #2 for generating PRS (see above) are subsets of approach #3, we conclude that their corresponding Type I error rates for PRSxE testing are also preserved.

**Table S1:** Estimated type I error and interaction effect for PRSxE using different PRS constructions methods.

| **PRS SNPs** | **Type I Error\***<br>**PRS x E** | **Estimated**<br>**$\beta_{PRS \times E}$** |
| --- | --- | --- |
| GWAS significant SNPs under threshold: |  |  |
| 5 x 10<sup>-5</sup> | 0.054 | -0.0036 |
| 5 x 10<sup>-3</sup> | 0.038 | -0.0049 |
| 5 x 10<sup>-1</sup> | 0.056 | -0.0033 |
| 20 main effects SNPs | 0.050 | -0.0029 |
| 10 main effects SNPs and 10 no-effects SNPs | 0.050 | 0.0033 |

\* Estimated Type I error and interaction effect based on 1,000 simulated replicates. Each replicate includes 20 SNPs with only main effects and no interaction

### Data Application: Colorectal Cancer

We compare the above approaches in an analysis of GxE interactions for colorectal cancer (CRC). We use data from the Functionally Informed Gene-environment Interaction (FIGI) study, a consortium that includes 108,649 subjects (51,350 CRC cases and 57,299 controls) drawn from 45 contributing studies. We focus on E=regular use of aspirin/NSAIDs (denoted NSAIDs from hereon), an exposure that has been repeatedly shown to reduce the risk of CRC^17–19^. A total of 78,253 subjects (33,937 cases, 44,316 controls) have complete data on NSAIDs use and are included in the analyses. Additional details of the study sample and definition of exposure are provided in Drew et al.^19^

The most recent and largest GWAS of CRC identified 204 SNPs that reached genomewide significance^16^. We apply the approaches described above to assess evidence that the PRS constructed from these SNPs interacts with NSAIDs to affect CRC risk. The overall PRS was constructed by first applying logistic regression within the FIGI sample to the 204 GWAS SNPs, with adjustment for study, sex, age, and three ancestry PCs (approach #2 described above). The log-odds ratios (“betas”) estimated from this model were used as the weights [w] to construct a PRS_i_, i=1, …, N for each study subject. To construct pPRS, we first used AnnoQ which successfully annotated 189 of the 204 SNPs to 265 protein-coding genes. The remaining 15 SNPs were mapped to non-coding genes and are ignored in this analysis. Application of PANTHER annotated 66 of the 265 genes to a total of 50 pathways (Figure 1), with pathways for the remaining 199 genes not identified. Among the 50 pathways, four of them included more genes than expected by chance alone at a false discovery rate (FDR) of 5%, identified by a Fisher’s Exact test in PANTHER (Table 2). These included the TGF-β/ signaling pathway (raw p=5.8×10^−6^), Alzheimer disease presenilin pathway (p=5.8×10^−5^), Gonadotropin-releasing hormone receptor pathway (p=4.8×10^−5^), and Cadherin signaling pathway (p=1.38×10^−3^). A total of 30 of the 204 SNPs were annotated to genes in these pathways. Subsets of the above PRS weights were utilized to construct the corresponding four pPRS scores. Logistic regression was used to estimate and test interactions, with adjustment for study, sex, age, and three principal components of ancestry. For each of the four pPRS x E tests, we report p-values unadjusted for multiple comparisons, with the rationale that each pathway-based PRS was constructed in advance using auxiliary information. The subsets of genes in the TGF-β signaling (TGF-β) pathway and Gonadotropin-releasing hormone receptor (GRHR) pathways are highly overlapped (Figure 2), as are genes in the Cadherin signaling (CADH) and Alzheimer’s disease presenilin(ALZ) pathways (Figure 3). These overlaps lead to significant correlations between the computed pPRS scores for TGF-β and GRHR (R^2^=0.58) and for CADH and ALZ (R^2^=0.71). Given these overlaps, we also constructed two additional pPRS scores based on SNPs within the combined subsets of TGF-β/GRHR genes and CADH/ALZ genes, respectively, and used analogous logistic regression models to test the corresponding pPRS x NSAIDs interactions.

## Funding source Acknowledgments

Genetics and Epidemiology of Colorectal Cancer Consortium (GECCO): National Cancer Institute, National Institutes of Health, U.S. Department of Health and Human Services (R01 CA059045, R01 CA201407, R01 CA273198). Genotyping/Sequencing services were provided by the Center for Inherited Disease Research (CIDR) contract number HHSN268201700006I and HHSN268201200008I. This research was funded in part through the NIH/NCI Cancer Center Support Grant P30 CA015704. Scientific Computing Infrastructure at Fred Hutch funded by ORIP grant S10OD028685.

The ATBC Study is supported by the Intramural Research Program of the U.S. National Cancer Institute, National Institutes of Health, Department of Health and Human Services.

CLUE II funding was from the National Cancer Institute (U01 CA086308, Early Detection Research Network; P30 CA006973), National Institute on Aging (U01 AG018033), and the American Institute for Cancer Research. The content of this publication does not necessarily reflect the views or policies of the Department of Health and Human Services, nor does mention of trade names, commercial products, or organizations imply endorsement by the US government. Maryland Cancer Registry (MCR) Cancer data was provided by the Maryland Cancer Registry, Center for Cancer Prevention and Control, Maryland Department of Health, with funding from the State of Maryland and the Maryland Cigarette Restitution Fund. The collection and availability of cancer registry data is also supported by the Cooperative Agreement NU58DP006333, funded by the Centers for Disease Control and Prevention. Its contents are solely the responsibility of the authors and do not necessarily represent the official views of the Centers for Disease Control and Prevention or the Department of Health and Human Services.

ColoCare: This work was supported by the National Institutes of Health (grant numbers R01 CA189184 (Li/Ulrich), U01 CA206110 (Ulrich/Li/Siegel/Figueiredo/Colditz, 2P30CA015704-40 (Gilliland), R01 CA207371 (Ulrich/Li)), the Matthias Lackas-Foundation, the German Consortium for Translational Cancer Research, and the EU TRANSCAN initiative.

The Colon Cancer Family Registry (CCFR, www.coloncfr.org) is supported in part by funding from the National Cancer Institute (NCI), National Institutes of Health (NIH) (award U01 CA167551). Support for case ascertainment was provided in part from the Surveillance, Epidemiology, and End Results (SEER) Program and the following U.S. state cancer registries: AZ, CO, MN, NC, NH; and by the Victoria Cancer Registry (Australia) and Ontario Cancer Registry (Canada). The CCFR Set-1 (Illumina 1M/1M-Duo) and Set-2 (Illumina Omni1-Quad) scans were supported by NIH awards U01 CA122839 and R01 CA143237 (to GC). The CCFR Set-3 (Affymetrix Axiom CORECT Set array) was supported by NIH award U19 CA148107 and R01 CA81488 (to SBG). The CCFR Set-4 (Illumina OncoArray 600K SNP array) was supported by NIH award U19 CA148107 (to SBG) and by the Center for Inherited Disease Research (CIDR), which is funded by the NIH to the Johns Hopkins University, contract number HHSN268201200008I. Additional funding for the OFCCR/ARCTIC was through award GL201-043 from the Ontario Research Fund (to BWZ), award 112746 from the Canadian Institutes of Health Research (to TJH), through a Cancer Risk Evaluation (CaRE) Program grant from the Canadian Cancer Society (to SG), and through generous support from the Ontario Ministry of Research and Innovation. The SFCCR Illumina HumanCytoSNP array was supported in part through NCI/NIH awards U01/U24 CA074794 and R01 CA076366 (to PAN). The content of this manuscript does not necessarily reflect the views or policies of the NCI, NIH or any of the collaborating centers in the Colon Cancer Family Registry (CCFR), nor does mention of trade names, commercial products, or organizations imply endorsement by the US Government, any cancer registry, or the CCFR.

COLO2&3: National Institutes of Health (R01 CA060987).

CPS-II: The American Cancer Society funds the creation, maintenance, and updating of the Cancer Prevention Study-II (CPS-II) cohort. The study protocol was approved by the institutional review boards of Emory University, and those of participating registries as required.

CRCGEN: Colorectal Cancer Genetics & Genomics, Spanish study was supported by Instituto de Salud Carlos III, co-funded by FEDER funds –a way to build Europe– (grants PI14-613 and PI09-1286), Agency for Management of University and Research Grants (AGAUR) of the Catalan Government (grant 2017SGR723), Junta de Castilla y León (grant LE22A10-2), the Spanish Association Against Cancer (AECC) Scientific Foundation grant GCTRA18022MORE and the Consortium for Biomedical Research in Epidemiology and Public Health (CIBERESP), action Genrisk. Sample collection of this work was supported by the Xarxa de Bancs de Tumors de Catalunya sponsored by Pla Director d’Oncología de Catalunya (XBTC), Plataforma Biobancos PT13/0010/0013 and ICOBIOBANC, sponsored by the Catalan Institute of Oncology. We thank CERCA Programme, Generalitat de Catalunya for institutional support.

DACHS: This work was supported by the German Research Council (BR 1704/6-1, BR 1704/6-3, BR 1704/6-4, CH 117/1-1, HO 5117/2-1, HE 5998/2-1, KL 2354/3-1, RO 2270/8-1 and BR 1704/17-1), the Interdisciplinary Research Program of the National Center for Tumor Diseases (NCT), Germany, and the German Federal Ministry of Education and Research (01KH0404, 01ER0814, 01ER0815, 01ER1505A and 01ER1505B).

DALS: National Institutes of Health (R01 CA048998 to M. L. Slattery).

EDRN: This work is funded and supported by the NCI, EDRN Grant (U01-CA152753).

Harvard cohorts: HPFS is supported by the National Institutes of Health (P01 CA055075, UM1 CA167552, U01 CA167552, R01 CA137178, R01 CA151993, and R35 CA197735), NHS by the National Institutes of Health (P01 CA087969, UM1 CA186107, R01 CA137178, R01 CA151993, and R35 CA197735), and PHS by the National Institutes of Health (R01 CA042182).

Hawaii Adenoma Study: NCI grants R01 CA072520.

Kentucky: This work was supported by the following grant support: Clinical Investigator Award from Damon Runyon Cancer Research Foundation (CI-8); NCI R01CA136726.

LCCS: The Leeds Colorectal Cancer Study was funded by the Food Standards Agency and Cancer Research UK Programme Award (C588/A19167).

MCCS: Melbourne Collaborative Cohort Study (MCCS) cohort recruitment was funded by VicHealth and Cancer Council Victoria. The MCCS was further augmented by Australian National Health and Medical Research Council grants 209057, 396414 and 1074383 and by infrastructure provided by Cancer Council Victoria. Cases and their vital status were ascertained through the Victorian Cancer Registry and the Australian Institute of Health and Welfare, including the Australian Cancer Database.

MEC: National Institutes of Health (R37 CA054281, P01 CA033619, and R01 CA063464).

MECC: This work was supported by the National Institutes of Health, U.S. Department of Health and Human Services (R01 CA081488, R01 CA197350, U19 CA148107, R01 CA242218, and a generous gift from Daniel and Maryann Fong.

NCCCS I & II: We acknowledge funding support for this project from the National Institutes of Health, R01 CA066635 and P30 DK034987.

NFCCR: This work was supported by an Interdisciplinary Health Research Team award from the Canadian Institutes of Health Research (CRT 43821); the National Institutes of Health, U.S. Department of Health and Human Services (U01 CA074783); and National Cancer Institute of Canada grants (18223 and 18226). The authors wish to acknowledge the contribution of Alexandre Belisle and the genotyping team of the McGill University and Génome Québec Innovation Centre, Montréal, Canada, for genotyping the Sequenom panel in the NFCCR samples. Funding was provided to Michael O. Woods by the Canadian Cancer Society Research Institute.

PLCO: Intramural Research Program of the Division of Cancer Epidemiology and Genetics and supported by contracts from the Division of Cancer Prevention, National Cancer Institute, NIH, DHHS. Funding was provided by National Institutes of Health (NIH), Genes, Environment and Health Initiative (GEI) Z01 CP 010200, NIH U01 HG004446, and NIH GEI U01 HG 004438.

SELECT: Research reported in this publication was supported in part by the National Cancer Institute of the National Institutes of Health under Award Numbers U10 CA037429 (CD Blanke), and UM1 CA182883 (CM Tangen/IM Thompson). The content is solely the responsibility of the authors and does not necessarily represent the official views of the National Institutes of Health.

SMS and REACH: This work was supported by the National Cancer Institute (grant P01 CA074184 to J.D.P. and P.A.N., grants R01 CA097325, R03 CA153323, and K05 CA152715 to P.A.N., and the National Center for Advancing Translational Sciences at the National Institutes of Health (grant KL2 TR000421 to A.N.B.-H.)

The Swedish Low-risk Colorectal Cancer Study (SLRCCS): The study was supported by grants from the Swedish research council; K2015-55X-22674-01-4, K2008-55X-20157-03-3, K2006-72X-20157-01-2 and the Stockholm County Council (ALF project).

UK Biobank: This research has been conducted using the UK Biobank Resource under Application Number 8614

VITAL: National Institutes of Health (K05 CA154337).

WHI: The WHI program is funded by the National Heart, Lung, and Blood Institute, National Institutes of Health, U.S. Department of Health and Human Services through contracts 75N92021D00001, 75N92021D00002, 75N92021D00003, 75N92021D00004, 75N92021D00005

Individual authors report the following funding support: Andrew Chan: R35 CA253185; Temitope Keku: U01 CA093326, R01 CA066635; Victor Moreno: Spanish Association Against Cancer (AECC) Scientific Foundation grant GCTRA18022MORE. Consortium for Biomedical Research in Epidemiology and Public Health (CIBERESP), action Genrisk

## Study-specific Acknowledgements

CCFR: The Colon CFR graciously thanks the generous contributions of their study participants, dedication of study staff, and the financial support from the U.S. National Cancer Institute, without which this important registry would not exist. The authors would like to thank the study participants and staff of the Seattle Colon Cancer Family Registry and the Hormones and Colon Cancer study (CORE Studies).

CLUE II: We thank the participants of Clue II and appreciate the continued efforts of the staff at the Johns Hopkins George W. Comstock Center for Public Health Research and Prevention in the conduct of the Clue II Cohort Study. Cancer data was provided by the Maryland Cancer Registry, Center for Cancer Prevention and Control, Maryland Department of Health, with funding from the State of Maryland and the Maryland Cigarette Restitution Fund. The collection and availability of cancer registry data is also supported by the Cooperative Agreement NU58DP006333, funded by the Centers for Disease Control and Prevention. Its contents are solely the responsibility of the authors and do not necessarily represent the official views of the Centers for Disease Control and Prevention or the Department of Health and Human Services.

CPS-II: The authors express sincere appreciation to all Cancer Prevention Study-II participants, and to each member of the study and biospecimen management group. The authors would like to acknowledge the contribution to this study from central cancer registries supported through the Centers for Disease Control and Prevention’s National Program of Cancer Registries and cancer registries supported by the National Cancer Institute’s Surveillance Epidemiology and End Results Program. The authors assume full responsibility for all analyses and interpretation of results. The views expressed here are those of the authors and do not necessarily represent the American Cancer Society or the American Cancer Society – Cancer Action Network.

DACHS: We thank all participants and cooperating clinicians, and everyone who provided excellent technical assistance.

EDRN: We acknowledge all contributors to the development of the resource at the University of Pittsburgh School of Medicine, Division of Gastroenterology, Hepatology and Nutrition, Department of Pathology, and Biomedical Informatics.

Harvard cohorts: The study protocol was approved by the institutional review boards of the Brigham and Women’s Hospital and Harvard T.H. Chan School of Public Health, and those of participating registries as required. We acknowledge Channing Division of Network Medicine, Department of Medicine, Brigham and Women’s Hospital as home of the NHS. The authors would like to acknowledge the contribution to this study from central cancer registries supported through the Centers for Disease Control and Prevention’s National Program of Cancer Registries (NPCR) and/or the National Cancer Institute’s Surveillance, Epidemiology, and End Results (SEER) Program. Central registries may also be supported by state agencies, universities, and cancer centers. Participating central cancer registries include the following: Alabama, Alaska, Arizona, Arkansas, California, Colorado, Connecticut, Delaware, Florida, Georgia, Hawaii, Idaho, Indiana, Iowa, Kentucky, Louisiana, Massachusetts, Maine, Maryland, Michigan, Mississippi, Montana, Nebraska, Nevada, New Hampshire, New Jersey, New Mexico, New York, North Carolina, North Dakota, Ohio, Oklahoma, Oregon, Pennsylvania, Puerto Rico, Rhode Island, Seattle SEER Registry, South Carolina, Tennessee, Texas, Utah, Virginia, West Virginia, Wyoming. The authors assume full responsibility for analyses and interpretation of these data.

Kentucky: We would like to acknowledge the staff at the Kentucky Cancer Registry.

LCCS: We acknowledge the contributions of all who conducted this study which was originally reported as 10.1093/carcin/24.2.275.

NCCCS I & II: We would like to thank the study participants, and the NC Colorectal Cancer Study staff.

PLCO: The authors thank the PLCO Cancer Screening Trial screening center investigators and the staff from Information Management Services Inc and Westat Inc. Most importantly, we thank the study participants for their contributions that made this study possible. Cancer incidence data have been provided by the District of Columbia Cancer Registry, Georgia Cancer Registry, Hawaii Cancer Registry, Minnesota Cancer Surveillance System, Missouri Cancer Registry, Nevada Central Cancer Registry, Pennsylvania Cancer Registry, Texas Cancer Registry, Virginia Cancer Registry, and Wisconsin Cancer Reporting System. All are supported in part by funds from the Center for Disease Control and Prevention, National Program for Central Registries, local states or by the National Cancer Institute, Surveillance, Epidemiology, and End Results program. The results reported here and the conclusions derived are the sole responsibility of the authors.

SELECT: We thank the research and clinical staff at the sites that participated on SELECT study, without whom the trial would not have been successful. We are also grateful to the 35,533 dedicated men who participated in SELECT.

SLRCCS: We would like to thank Annika Lindblom and the SLRCCS Study staff and participants.

WHI: The authors thank the WHI investigators and staff for their dedication, and the study participants for making the program possible. A full listing of WHI investigators can be found at: https://s3-us-west-2.amazonaws.com/www-whi-org/wp-content/uploads/WHI-Investigator-Long-List.pdf

